# Rapid evolution of antiviral *APOBEC3* genes driven by the conflicts with ancient retroviruses

**DOI:** 10.1101/707190

**Authors:** Jumpei Ito, Robert J. Gifford, Kei Sato

## Abstract

The evolution of antiviral genes has been fundamentally shaped by antagonistic interactions with ancestral viruses. The *AID/APOBEC* family genes (*AID* and *APOBEC1-4*) encode cellular cytosine deaminases that target nucleic acids and catalyze C-to-U mutations. In the case of retroviral replication, APOBEC3 proteins induce C-to-U mutations in minus-stranded viral DNA, which results in G-to-A mutations in the viral genome. Previous studies have indicated that the expansion and rapid evolution of mammalian *APOBEC3* genes has been driven by an arms race with retroviral parasites, but this has not been thoroughly investigated. Endogenous retroviruses (ERVs) are retrotransposons originated from ancient retroviral infections. These sequences sometimes bear the hallmarks of APOBEC3-mediated mutations, and therefore serve as a record of the ancient conflict between retroviruses and *APOBEC3* genes. Here we systematically investigated the sequences of ERVs and *APOBEC3* genes in mammals to reconstruct details of the evolutionary conflict between them. We identified 1,420 *AID/APOBEC* family genes in a comprehensive screen of mammalian genome. Of the *AID/APOBEC* family genes, *APOBEC3* genes have been selectively amplified in mammalian genomes and disclose evidence of strong positive selection - whereas the catalytic domain was highly conserved across species, the structure loop 7, which recognizes viral DNA/RNA substrates, was shown to be evolving under strong positive selection. Although *APOBEC3* genes have been amplified by tandem gene duplication in most mammalian lineages, the retrotransposition-mediated gene amplification was found in several mammals including New World monkeys and prosimian primates. Comparative analysis revealed that G-to-A mutations are accumulated in ERVs, and that the G-to-A mutation signatures on ERVs is concordant with the target preferences of APOBEC3 proteins. Importantly, the number of *APOBEC3* genes was significantly correlated with the frequency of G-to-A mutations in ERVs, suggesting that the amplification of *APOBEC3* genes led to stronger attacks on ERVs and/or their ancestral retroviruses by *APOBEC3* proteins. Furthermore, the numbers of *APOBEC3* genes and ERVs in mammalian genomes were positively correlated, and in primates, the timings of *APOBEC3* gene amplification was concordant with that of ERV invasions. Our findings suggest that conflict with ancient retroviruses was a major selective pressure driving the rapid evolution of *APOBEC3* genes in mammals.

## Introduction

Gene evolution is driven by a variety of factors including interactions with pathogens^1,2^. For instance, the genes encoding major histocompatibility complex have diversified and evolved to combat various pathogens as a central player of acquired immunity in mammals^3^. These insights indicate that the evolutionary conflicts between hosts and their invaders have shaped the evolution of genes that are important for protecting the hosts from pathogens.

Activation-induced cytidine deaminase/apolipoprotein B mRNA editing enzyme, catalytic polypeptide-like (AID/APOBEC) superfamily proteins are cellular cytosine deaminases that catalyze C-to-U mutations in nucleotides. AID/APOBEC family proteins contain conserved zinc-dependent catalytic domain (Z domain) with the HxE/SPCxxC motif and are closely associated with important phenomena found in vertebrates such as immunity, malignancy, metabolism, and infectious diseases (reviewed in refs. ^4,5^). For instance, AID induces somatic hypermutation in B cells and promotes antibody diversification^5^, and APOBEC1 (A1) regulates lipid metabolism by enzymatically editing the mRNA of apolipoprotein B gene^6^. Although the physiological roles of APOBEC2 (A2) and APOBEC4 (A4) remain unknown, it is known that APOBEC3 (A3) proteins play a pivotal role in restricting retroviral replication (reviewed in ref. ^7^).

Mammalian A3 proteins are antiviral factors that restrict the replication of retroviruses^7^ and non-retroviruses^8–10^. The evolutionary conflict between human A3G protein and human immunodeficiency virus type 1 (HIV-1) has been studied particularly intensively. Human A3G proteins are incorporated into HIV-1 particle and enzymatically induce C-to-U mutations in viral complementary DNA, which results in G-to-A mutations in viral genome^11,12^. Although A3G-mediated mutations lead to the accumulation of lethal and missense mutations in viral genome and ultimately abolish viral replication, an HIV-1-encoding protein, viral infectivity factor (Vif), counteracts this antiviral action by degrading A3G via a ubiquitin-proteasome-dependent manner (reviewed in ref. ^7^). Such conflicts between A3 proteins and modern viruses (particularly retroviruses) have been reported in a broad range of mammalian species and viruses infecting with them (reviewed in ref. ^13^).

Endogenous retroviruses (ERVs) are retrotransposon lineages that are thought to have originated from ancient exogenous retroviruses via infection of germline cells^14,15^. ERVs currently occupy a substantial fraction of mammalian genomes, suggesting that a massive amount of ERVs have invaded the host genomes. To combat such genomic invasions by retroviral parasites, the hosts have developed defense systems such as KRAB zinc-finger proteins^16^ and PIWI-interacting RNAs^17^. Interestingly, previous studies reported the suppressive effects of A3 proteins on ERVs and/or their ancestral exogenous retroviruses. A3 proteins can suppress the replications of ERVs in cell cultures^14,18^ and in a transgenic mouse model^19^. Furthermore, previous studies addressed the G-to-A hypermutation signature on ERVs in various mammals and proposed that this signature is a sign of the ancient attack by A3 proteins on ERVs and/or their ancestral retroviruses^14,15,18,20^. Together, these insights suggest a long-standing evolutionary conflict between A3 proteins and retroviruses including ERVs.

Although the *AID/APOBEC* family genes except for *A3* are conserved in various vertebrates, *A3* genes are specifically encoded in placental mammals^4^. Moreover, although *AID, A1, A2* and *A4* genes are singly encoded in each vertebrate including mammals, *A3* genes are under positive selection^21–23^ and are highly amplified in multiple mammalian lineages such as primates^24^. These insights suggest that *A3* genes had been acquired and have rapidly evolved and diversified during mammalian evolution. Based on the sequences of their Z domains, *A3* genes are classified into three classes, *A3Z1, A3Z2* and, *A3Z3* (reviewed in refs. ^7,13,24^). For example, human *A3* genes are composed of 7 paralogs (A3A, *A3B, A3C, A3D, A3F, A3G*, and *A3H*). Of these genes, *A3A, A3C* and *A3H* (also known as *A3Z1, A3Z2* and *A3Z3* in the other mammals) respectively contain single Z domain, while the other four genes harbor double Z domains: *A3Z2-A3Z1* for *A3B* and A3G, *A3Z2-A3Z2* for *A3D* and *A3F*^13,24^. The driving forces behind the amplification and diversification of *A3* genes remain unclear. However, an attractive hypothesis is that it has been driven by conflict between host species and retroviruses. In this study, we examine the evolutionary history of ERVs and *A3* genes in the genomes of 160 mammalian species. We newly identify more than 1,000 sequences of *A3* Z domains in mammalian genome and uncover a hidden diversity of mammalian *A3* genes. Furthermore, we show the evidences suggesting that the rapid evolution of *A3* genes, including gene amplification and diversification, was triggered by retroviral invasions and that the amplification of *A3* genes led to stronger attacks on retroviruses by A3 proteins. To our knowledge, this is the first study comprehensively addressing the evolutionary conflict of ERVs and mammalian *A3* genes.

## Results

### Identification and classification of mammalian *AID/APOBEC* family genes

We screened the genome sequences of 160 mammalian species (**Supplementary Table 1**) and extracted sequences homologous to Z domains of *AID/APOBEC* family genes using a tBLASTn-based method (http://giffordlabcvr.github.io/DIGS-tool/). Amino acid sequences of *AID/APOBEC* family genes of five mammals (human, mouse, cow, megabat, and cat) were used as queries of the search (**Supplementary Table 2** and **Supplementary Fig. 1**). Of the extracted sequences, we particularly focused on the sequence region corresponding to the conserved sequence of Z domains of *AID/APOBEC* family genes (**Supplementary Fig. 1a** and **1c**)^24^. We identified a total of 1,420 distinct loci disclosing homology to Z domains of *AID/APOBEC* family genes (**Supplementary Table 3**). Phylogenetic reconstructions revealed that these loci group into nine clades, of which seven are canonical Z domains of *AID/APOBEC* family genes (*AID, A1, A2, A3Z1, A3Z2, A3Z3*, and A4) (**Fig. 1a** and **1b**). We also identified additional, previously uncharacterized groups, designated unclassified *AID/APOBEC* genes 1 (*UA1*) and 2 (*UA2*) (**Fig. 1a** and **1b**). *UA1* genes were specifically detected in the basal mammalian lineages Afrotheria (elephants, tenrecs, and sea cows) and Xenarthra (armadillos), while *UA2* genes were encoded in marsupials (infraclass Marsupialia) (**Fig. 1c**). These phylogenetic relationships were supported by multiple methods (**Fig. 1a** and **Supplementary Fig. 2**), and HxE and SPCxxC motifs corresponding to the canonical catalytic domain of AID/APOBEC proteins were found in *UA1* and *UA2* gene sequences (**Supplementary Fig. 3a**). Furthermore, *UA1* and *UA2* genes were under the purifying selection (**Supplementary Fig. 3b**). These suggest that *UA1* and *UA2* genes are protein-coding sequences and part of the *AID/APOBEC* family. In fact, the *UA2* gene in opossum (*Monodelphis domestica*) was annotated as *‘APOBEC5’* in a previous study^25^.

**Fig. 1.**
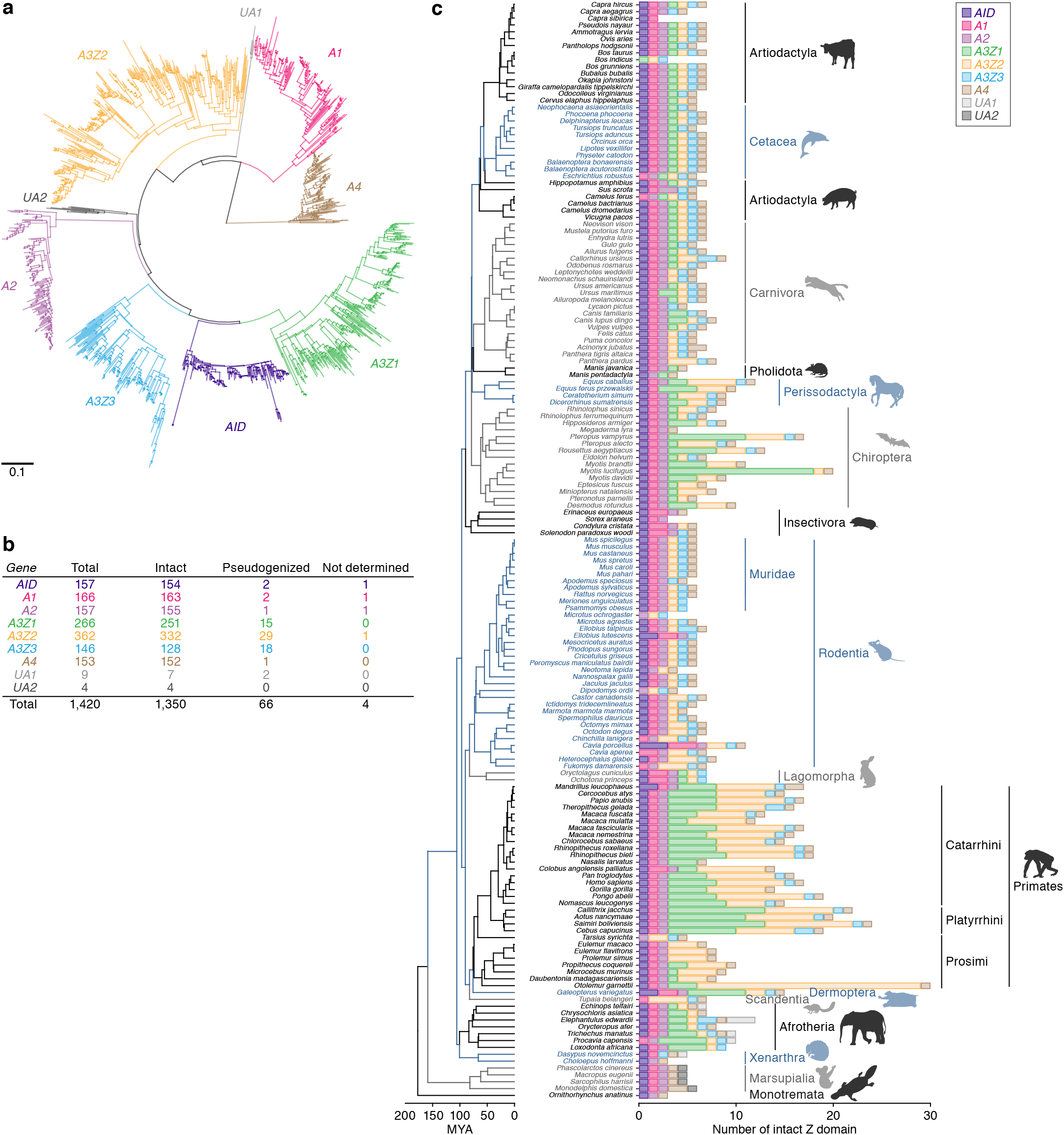
Identification of the *AID/APOBEC* Z domains in mammalian genomes. **a** Phylogenetic tree of the identified sequences of *AID/APOBEC* Z domains in 160 mammalian genomes. The tree was constructed using neighbor-joining (NJ) method^66^ based on the nucleic acid sequences. Scale bar indicates the genetic distance. **b** Number of the identified *AID/APOBEC* Z domains. “Intact”, Z domain without premature stop codons in the conserved sequence; “pseudogenized”, Z domain with the stop codons in the sequence; “not determined”, Z domain with ambiguous nucleotides in the sequence. **c** Number of the identified *AID/APOBEC* Z domains in each mammal species. The count of intact Z domains is only indicted. The count including both of the intact and pseudogenized Z domains is shown in **Supplementary Fig. 4**. Species tree is derived from TimeTree database^77^.

As summarized in **Fig. 1b** we detected 157 *AID*, 166 *A1*, 157 *A2*, 266 *A3Z1*, 362 *A3Z2*, 146 *A3Z3*, 153 *A4*, 9 *UA1*, and 4 *UA2* genes in 160 species of mammalian genomes. Interestingly, *A3Z1* and *A3Z2* genes were highly amplified, while the other family genes were not (**Fig. 1b** and **1c**). We also found that some sequences, particularly those of *A3* genes, were pseudogenized (**Fig. 1b**).

**Fig. 1c** and **Supplementary Fig. 4** illustrate the numbers of the Z domains of *AID/APOBEC* family genes identified in each mammalian species. The numbers of *A3* Z domains were different among species, and particularly, *A3Z1* and *A3Z2* genes in Perissodactyla, Chiroptera, Primates, and Afrotheria were highly amplified. Consistent with previous reports^21,26,27^, canonical *A3* genes were not detected in marsupials and Monotremata. Also, *A3Z1* was commonly absent in Rodentia, and *A3Z3* was absent in Strepsirrhini and Microchiroptera. Higher amplifications of *A3Z3* gene were not detected in any mammals. Although the duplication of *A3Z3* gene was specifically observed in Carnivora, the duplicated ones were pseudogenized (**Supplementary Fig. 5**).

### Evolutionary history of mammalian *A3* genes

To investigate the evolutionary history of mammalian *A3* genes, we examined sequence characteristics of *AID/APOBEC* genes. As shown in **Fig. 2a**, the positional conservation scores (Shannon entropy scores) in *A3Z1, A3Z2*, and *A3Z3* genes tended to be much higher than those of the other *AID/APOBEC* family genes, indicating strong positive selection favoring diversification. We then detected the codon sites under the positive selection through dN/dS ratio test (branch-site model^28^). Although the catalytic domains, which are composed of HxE and SPCxxC motifs^4,5,7^, were highly conserved in the seven AID/APOBEC family proteins, we detected substantial numbers of positively selected sites in A3 proteins (**Fig. 2b**). Notably, comparisons to the amino acid positions to those of human A3A (A3Z1 ortholog in primates)^29^, A3C (A3Z2 ortholog in primates)^30^, and A3H (A3Z3 ortholog in primates)^31^ revealed that the positively selected sites are preferentially detected in a structural region, called loop 7, which recognizes substrate nucleic acids (**Fig. 2b**). Moreover, most of the positively selected sites are located on the protein surface (**Fig. 2b**).

**Fig. 2.**
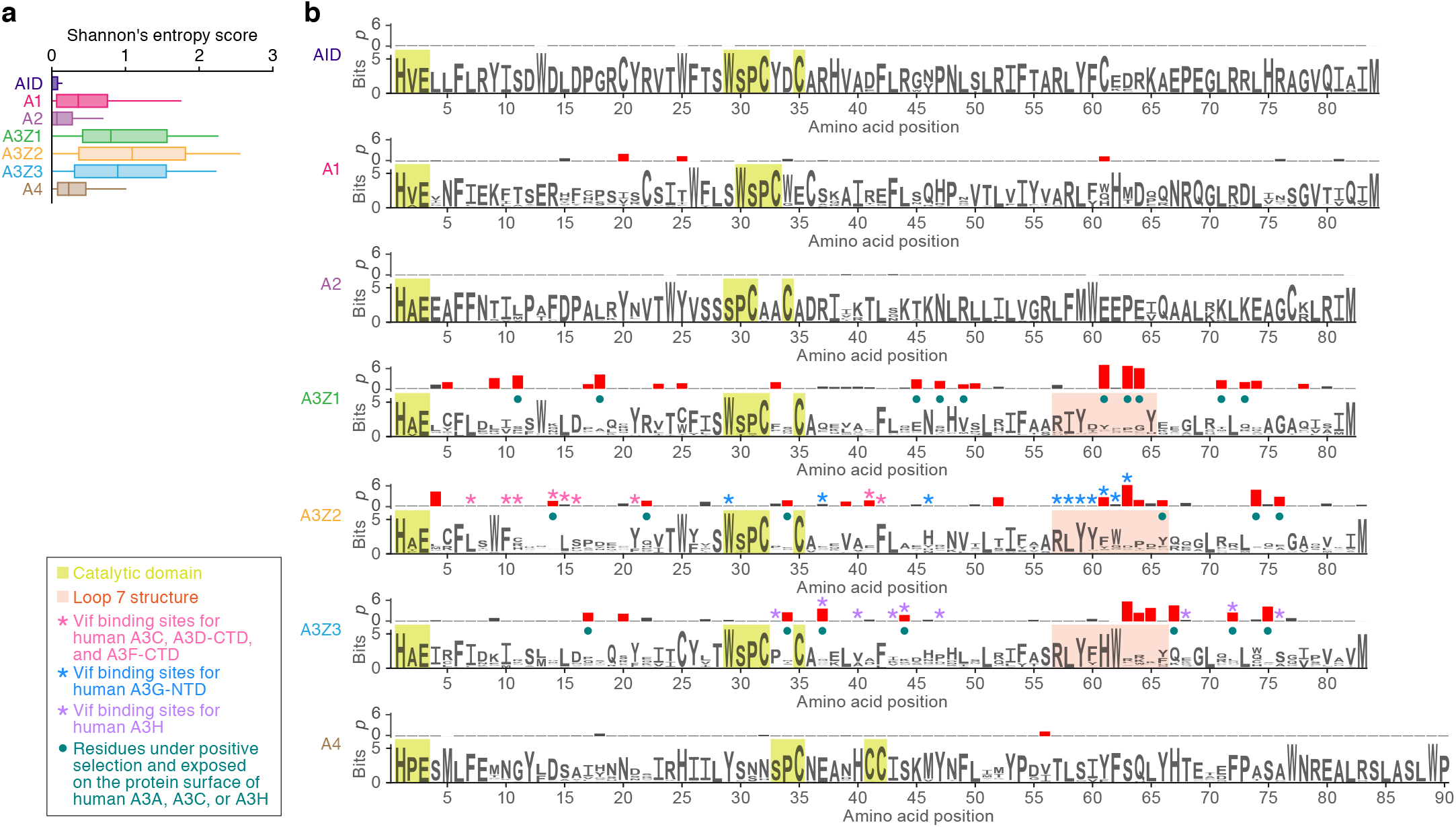
Evolutionary features of AID/APOBEC Z domains. **a** Difference in the sequence conservations among seven classes of AID/APOBEC Z domains. Positional sequence conservation scores (Shannon’s entropy scores) were calculated in respective amino acid sites of the multiple sequence alignment (shown as logo plots in **b**). **b** (Bottom) Logo plots of the conserved sequences of the AID/APOBEC Z domains. Yellow square indicates the amino acid residues comprising the catalytic domain of AID/APOBEC proteins. Pink square indicates the amino acid residues corresponding to the structure loop 7. The other characteristics on each amino acid residue are summarized in the box at the lower left of the panel. (Top) The *p* values (-log10) in dN/dS ratio test (with branch-site model^28^) at each codon site. The sites under the positive selection with statistically significance (*p* <0.05) are indicated by red bars.

Investigation of amplified *A3* loci revealed that the majority of *A3* genes are encoded in the canonical *A3* genomic locus, which is sandwiched by *CBX6* and *CBX7* genes (**Fig. 3a** and **Supplementary Table 4**), as previously described^13,24^. These results suggest that the amplifications of *A3* genes were mainly occurred as the tandem gene duplication. However, exceptions to this rule were identified in primates: the genomes of three species, *Saimiri boliviensis, Aotus nancymaae*, and *Otolemur garnettii*, were found to encode more *A3* loci outside the canonical locus than within it (**Fig. 3b**). When we assessed the genomic loci of the *A3* genes in these three primates, most of the *A3* genes were encoded at entirely distinct loci each other (**Fig. 3c**). Furthermore, these *A3* genes in New World monkeys (*Saimiri boliviensis* and *Aotus nancymaae*) exhibit intron-less structures (**Supplementary Fig. 6a**), which would have been originated via mRNA splicing followed by reverse transcription^32^. These findings suggest that the *A3* genes encoded at the outside of the canonical *A3* locus likely originated via retrotransposition. These retrotransposed *A3* genes in New World monkeys were more closely related to human *A3G* gene compared to the other double-domain *A3* genes in human (**Supplementary Fig. 6b**). Although most of these retrotransposed *A3* genes are pseudogenized (**Fig. 3d**), some of them retain relatively longer open reading frames (ORFs) (**Supplementary Fig. 6c**). In particular, one of these retrotransposed *A3* genes in *Aotus nancymaae* (referred to as “outside #3”) retains a full-length ORF (**Supplementary Fig. 6c**). Moreover, we analyzed public RNA-sequencing (RNA-Seq) data and revealed that mRNA of this outside #3 was expressed in a broad range of tissues of *Aotus nancymaae* (**Supplementary Fig. 7**). Further, we found that this outside #3 is annotated as *A3G* gene of *Aotus nancymaae* (ENSANAG00000031271) in the Ensembl gene database (http://www.ensembl.org; Release 97). Taken together, A3G-like genes were amplified via retrotransposition in New World monkeys, and some of these amplified genes seem to be functional.

**Fig. 3.**
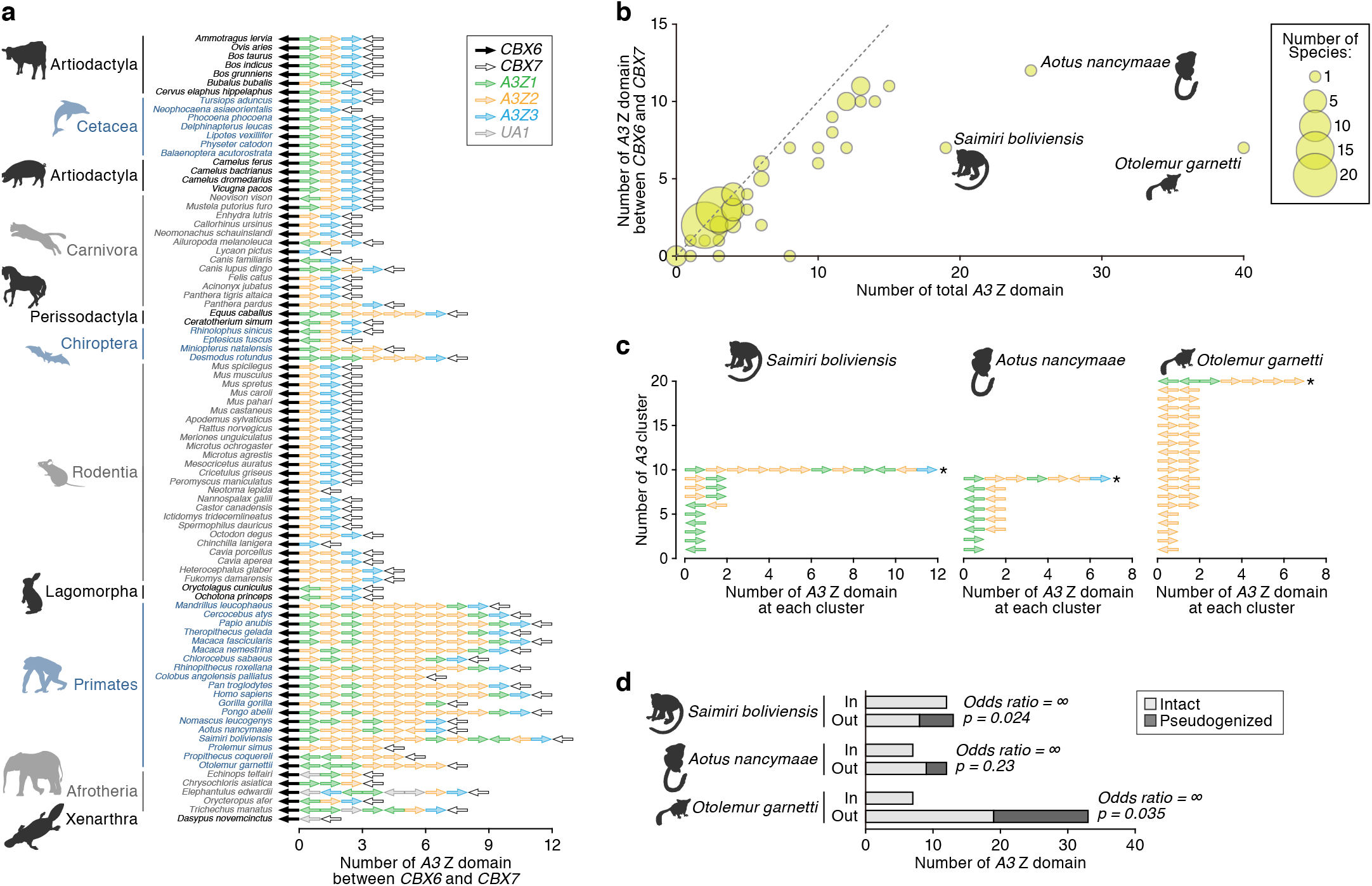
Genomic location of *A3* genes. **a** Genomic order of the *AID/APOBEC* Z domains within the canonical *A3* gene locus, which is sandwiched by *CBX6* and *CBX7* genes. Mammalian genomes in which *CBX6* and *CBX7* genes were detected in the same scaffold were only analyzed. The arrows indicate the direction of respective loci. **b** Bubble plot of the number of *A3* Z domains in mammals. The number of the *A3* Z domains in the whole genome (x-axis) and that within the canonical *A3* gene locus (y-axis) in each mammal are plotted. Dot size is proportional to the number of species. **c** Genomic locations of *A3* Z domains in *Saimiri boliviensis, Aotus nancymaae*, and *Otolemur garnetti. A3* Z domains within 100 kb each other were clustered. An asterisk denotes the *A3* cluster corresponding to the canonical *A3* gene locus. The arrows indicate the direction of respective loci. **d** Association between the genomic location of *A3* genes and the pseudogenization. “In” or “Out” respectively denote the numbers of *A3* Z domains located inside or outside the canonical *A3* gene locus. Results for *Saimiri boliviensis, Aotus nancymaae*, and *Otolemur garnetti* are shown. Odds ratio and *p* value calculated with Fisher’s exact test are shown.

### Footprint of A3 activity in ERVs and its association with *A3* gene amplifications

To explore the impact of A3 activity on ERVs and their ancestor of ancient exogenous retroviruses, we performed comparative analysis of transposable elements (TEs) including ERVs in 160 species of mammals. As shown in **Fig. 4a** and **Supplementary Fig. 8**, the TE composition of mammalian species varies with respect to the proportions of DNA transposons, SINEs, LINEs, and ERVs. To investigate the accumulation level of G-to-A mutations in ERVs, we measured the strand bias of the G-to-A mutation rate in ERVs and other TEs. Since A3 protein induces G-to-A mutations selectively on the positive strand of retroviruses, the strand bias can be regarded as an indicator of the A3 attack on retroviruses. As shown in **Fig. 4b**, G-to-A mutations were accumulated preferentially on the positive strand of human ERVs but not of the other human TEs. We next classified mutation patterns based on the dinucleotide context. As shown in **Fig. 4c**, ERVs in the human genome preferentially exhibited either <u>G</u>G-to-<u>A</u>G or <u>G</u>A-to-<u>A</u>A mutations, consistent with the reported preferences of human A3G (<u>G</u>G-to-<u>A</u>G) and A3D, A3F, and A3H (<u>G</u>A-to-<u>A</u>A mutations)^11,33–39^. Additionally, some ERVs exhibited G-to-A hypermutations (**Fig. 4d**).

**Fig. 4.**
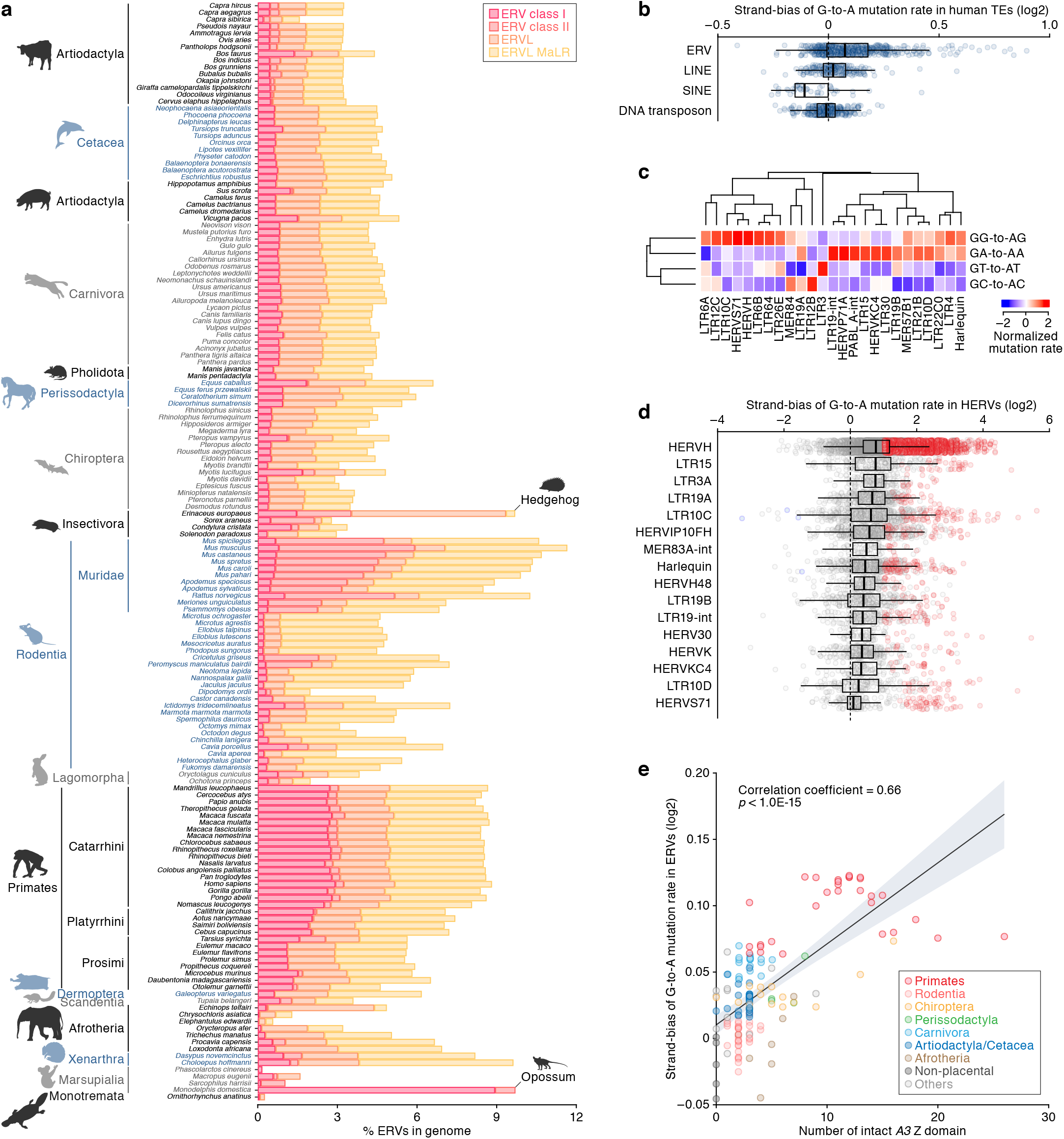
Footprint of the A3 activity in ERVs and its association with *A3* gene amplifications. **a** Proportions of ERV sequences in the genomes of respective mammalian species. Proportions of sequences of TEs (including LINE, SINE, and DNA transposon) in the genomes of respective mammalian species are also shown in **Supplementary Fig. 9**. **b** Strand bias scores of G-to-A mutation rates in human TEs (log2-transformed). The strand bias score is calculated as the G-to-A mutation rate ratio between the positive and negative strands (i.e., the mutation rate in the positive strand divided by the one in negative strand). That is, positive and negative values respectively indicate the positive and negative strand biases. Dots indicate the strand bias scores of respective TE groups. **c** Dinucleotide sequence composition of G-to-A mutation sites in human ERVs. Of the top 50 ERVs with respect to the strand bias score, the top 25 ERVs with respect to the variation among the four G-to-A mutation sites (<u>G</u>A, <u>G</u>T, <u>G</u>G, <u>G</u>C) are shown. **d** ERV copies presenting the G-to-A hypermutation signature. ERV copies with >1 log2-transformed strand bias score and <0.1 false discovery rate are indicated as red. **e** Association of the number of *A3* Z domains with the accumulation level of G-to-A mutations in ERVs in mammals. The x-axis indicates the number of intact *A3* Z domains, and the y-axis indicates the mean value of the log2-transformed strand bias scores among ERVs in the genome. Correlation coefficient and *P* value are calculated by Pearson’s correlation.

To explore the impact of the amplification of *A3* genes on ERVs, we assessed the accumulation level of G-to-A mutations in ERVs across mammals (**Supplementary Fig. 9**). Subsequently, we examined the association of the number of *A3* Z domains with the accumulation level of G-to-A mutations in ERVs and found the strong positive correlation (**Fig. 4e**) (Pearson’s correlation coefficient = 0.69, *p* < 1.0E-15). Collectively, these findings suggest that ERVs in mammals having no or fewer *A3* genes (e.g., non-placental mammals or rodents) exhibited lower accumulation levels of G-to-A mutations, whereas ERVs in mammals having a higher number of *A3* genes (e.g., Simiiformes and some chiropterans) exhibited higher accumulation levels of G-to-A mutations.

### Association between *A3* gene family expansion and ERV invasion

Finally, we examined the association between ERV invasions and *A3* gene family expansion. As shown in **Fig. 5a** and **5b**, we found that the number of *A3* Z domain was positively associated with the percentage of ERVs in mammalian genome (in Poisson regression, coefficient = 0.14, *p* <1.0E-15). Thus, species invaded by a larger amount of ERVs tended to have a higher number of *A3* genes. Exceptions occur in the rodent family Muridae, and two other species, hedgehog (*Erinaceus europaeus*), and opossum (*Monodelphis domestica);* all of these species have undergone a larger number of retroviral genome invasions but possessed few or no *A3* genes (**Supplementary Fig. 10**). As expected, ERVs in these species exhibited lower accumulation levels of G-to-A mutations (**Fig. 5b**). Furthermore, insertion dates of ERVs in these species were relatively young (**Supplementary Fig. 11** and **12**). Thus, ERVs seem to have prominently invaded in the genomes of these species without suffering from the intensive mutation-mediated attacks by A3 proteins.

**Fig. 5.**
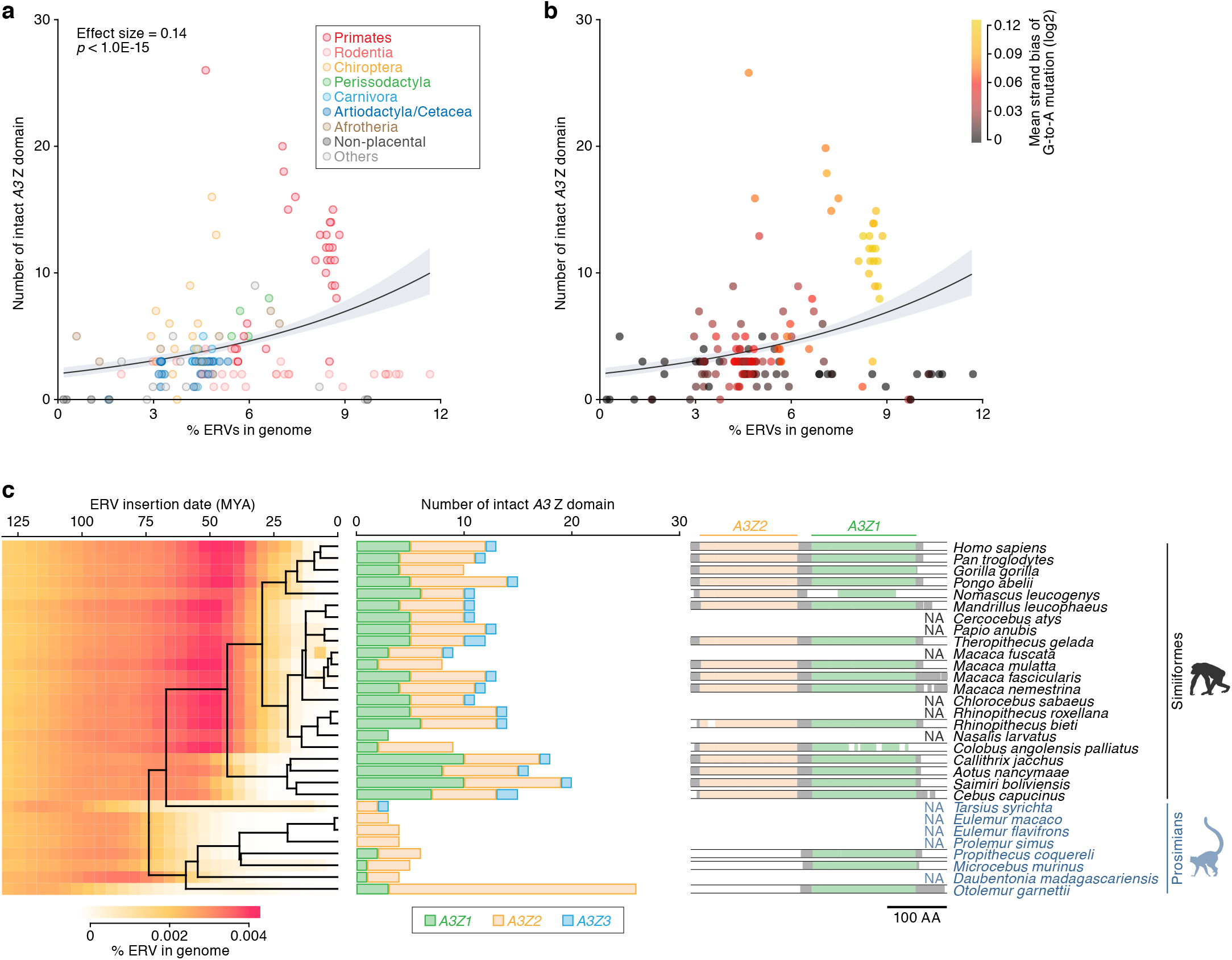
Association between *A3* gene family expansion and ERV invasion. **a, b** Association of the number of *A3* Z domains with the amount of ERV insertions in the genome. Dots are colored according to the species taxa (**a**) or the accumulation level of G-to-A mutations in ERVs (**b**). The association was evaluated under the Poisson regression with log link function. **c** Association of the timings between ERV invasion and the *A3* gene amplification in Primates. (Left) Amount of ERV insertions at each age in respective primate species. ERV insertion date was estimated based on the genetic distance of each ERV integrant from the consensus sequence under the molecular clock assumption (2.2 × 10^−9^ mutation/site/year^75^). (Middle) Number of intact *A3* Z domains. (Right) Schematic of the multiple sequence alignment of *A3G (A3Z2-Z3Z1* type) gene. Sequences of *A3G* genes in Primates recorded in the Ensembl gene database (http://www.ensembl.org) were used. NA, not applicable (no available data).

To deeply illuminate the association of *A3* gene family expansion with ERV invasion, we focused on the timing of these two events in Primates. Through assessing the age of ERV invasions in each species according to the genomic distance of each ERV integrant form the consensus sequence, we found that ERVs prominently invaded in the ancestors of Simiiformes (including Hominoidea, Old World monkeys, and New World monkeys) around 50 million years ago (MYA) (**Fig. 5c, left**). On the other hand, ancestors of Prosimians (including Lemurs, Lorisoids, and Tarsiers) have not experienced such prominent ERV invasion in that period. Furthermore, Simiiformes encoded higher numbers of *A3* genes, whereas Prosimians encoded lower numbers of *A3* genes (except for *Otolemur garnettii*), suggesting that the *A3* gene amplification occurred during early divergence of Simiiformes (**Fig. 5c, middle**). Moreover, we investigated the timing of the formation of the double-domain *A3G* gene (i.e., *A3G* gene with *A3Z2-A3Z1* structure) using the Ensembl gene database (www.ensembl.org/). We found that Simiiformes encoded the double-domain (*A3Z2-A3Z1) A3G* gene, whereas Prosimians did not, suggesting that the formation of the double-domain *A3G* gene also occurred during that period (**Fig. 5c, right**). Absence of the double-domain *A3G* gene in Prosimians is supported by the finding that no *A3Z2-A3Z1* genetic structures were observed in Prosimian genomes (**Fig. 3c**). Taken together, the timings of *A3* gene amplification and diversification were highly concordant to the timing of the prominent ERV invasions in Primates.

## Discussion

At present, mammalian A3 proteins are considered to represent antiviral factors against various retroviruses^13^ and non-retroviruses^8–10^. Although A3 family proteins are capable of targeting a broad range of viruses, the target range of individual A3 proteins may be relatively specific. For instance, human A3G can antagonize lentiviruses^12,40,41^ and gammaretroviruses^11,40,42^ including murine ERVs^43^ but is incapable of efficiently inhibiting betaretroviruses such as simian retrovirus^44^ and HERV-K^14,45^. On the other hand, human A3F potently inhibits the replication of betaretroviruses^14,44,45^. These observations suggest that the *A3* family genes have amplified and diversified to target a broad range of ancient viruses or ERVs; however, this has not been fully investigated. In the present study, we systematically characterized *A3* genes and ERVs in 160 mammalian species and shed light on such ancient conflicts between *A3* family genes and ERVs and/or their ancestral retroviruses.

We showed that species invaded by a larger amount of ERVs tended to encode more *A3* Z domains in the genomes (**Fig. 5a** and **5b**). We also demonstrated that the amplification of *A3* Z domains occurred around the same time to the prominent genomic invasion of ERVs or their ancestral retroviruses in primates (**Fig. 5c**). These findings suggest that such retroviral genomic invasion was one of the major driving forces for the amplification of mammalian *A3* genes. Furthermore, we demonstrated that the accumulation level of G-to-A mutations in ERVs was positively associated with the number of *A3* Z domains in the genomes (**Fig. 4e**). This finding strongly supports the concept proposed previously^14,15,18,20^ that the accumulation of G-to-A mutations in ERVs is due to A3 proteins. Moreover, this finding also suggests that the amplification of *A3* genes led to a stronger attack by A3 proteins on ERVs and/or their ancestral retroviruses. Taken together, our findings provide an evidence that diversification of mammalian *A3* genes has been shaped by the conflicts with retroviruses that were actively invading the genomes of mammalian ancestors.

We demonstrated that a structural region of A3 proteins, called loop 7, was under strong positive selection (**Fig. 2b**). Since loop 7 structures of A3 proteins determine the sequence specificity of viral nucleotide substrates^46^, our findings suggest that various ancestral retroviruses and ERVs could be a particular driver accelerating the rapid evolution of the *A3* genes encoding the loop 7 region. In addition, it is known that HIV-1 Vif specifically binds to the loop 7 structure of human A3G, leading to its degradation^47,48^. This raises a possibility that the Vif-like proteins encoded by ancestral retroviruses and/or ERVs could be another selective pressure on A3s, particularly loop 7 of the folded protein. In fact, all endogenous lentiviruses encode *vif* gene-like ORFs^49–51^. Therefore, it is reasonable to conclude that ancestral lentiviral Vif-like proteins have exerted selective pressure on mammalian *A3* genes, particularly on the loop 7 region. Consistent with this, our analysis shows that most of the sites under positive selection were located on the surface of A3 proteins (**Fig. 2b**). Moreover, it has been recently reported that Epstein-Barr virus (EBV) and related herpesviruses encode proteins that degrade human A3, and that these A3 antagonists specifically recognize loop 7^8^. These findings further suggest that other exogenous viruses could be another evolutionary pressure against mammalian A3s, particularly on the loop 7 structure.

The majority of *A3* genes are encoded in the canonical *A3* locus and have been amplified by tandem gene duplication (**Fig. 3a** and **3b**). However, we also detected the amplification of *A3* genes outside this region specifically in three primates (*Saimiri boliviensis, Aotus nancymaae*, and *Otolemur garnetti*) via retrotransposition (**Fig. 3b** and **3c**, and **Supplementary Fig. 6**). Particularly, A3G-like genes were amplified via retrotransposition in New World monkeys (*Saimiri boliviensis* and *Aotus nancymaae*), and some of these retrotransposed genes seem to be functional (**Supplementary Fig. 6** and **Fig. 7**). The creation of a novel anti-retroviral gene via retrotransposition in New World monkeys has been documented previously: the *TRIMCYP* fusion gene found in New World monkeys (genus *Aotus*) was created via distinct retrotransposition events^52,53^. This remarkable example of convergence suggests the strong selective pressure that retroviruses exert on this primate.

It is known that uncontrolled *A3* expression can be harmful for the hosts. In fact, exogenous expression of human A3A (A3Z1 ortholog) in cell cultures triggers cytotoxic effects^54–56^. Similarly, the aberrant expression of some human A3 proteins, particularly A3A^57,58^, A3B (A3Z2-A3Z1 ortholog)^57–60^, and A3H (A3Z3 ortholog)^61^, can contribute to cancer development by inducing somatic G-to-A mutations in human genome. In this study, we observed the gene loss as well as gene amplification of *A3* genes in multiple lineages of mammals (**Fig. 1c** and **Supplementary Fig. 4**). For example, *A3Z1* gene was lost in Rodentia, and *A3Z3* gene was lost in Strepsirrhini and Microchiroptera. These findings might be attributed to genotoxic potential of these *A3* genes. More interestingly, although *A3Z3* gene is highly conserved in most mammals except for some lineages (e.g., marsupials, microbats, and prosimians), *A3Z3* gene is not amplified in most mammalian lineages unlike *A3Z1* and *A3Z2* genes. As exceptions for *A3Z3* duplication, we detected the duplicated *A3Z3* genes specifically in carnivores and some species; however, almost all duplicated *A3Z3* genes were pseudogenized (**Supplementary Fig. 5**). Moreover, the duplication events happened twice independently during carnivore evolution. These observations strongly support the idea that *A3Z3* gene is indispensable for the hosts, but its duplication might be genotoxic. Taken together, our findings suggest the presence of the evolutionary constraints in *A3* gene amplification due to the genotoxic effects of A3 proteins.

Although A3 proteins can suppress retroviral replication in a G-to-A mutation-independent fashion (e.g., inhibition of reverse transcription)^62–65^, we here did not address this aspect because of the technical difficulty to assess the mutation-independent effect of A3 proteins on ERV invasions, based only on the genomic information. It should also be noted that the number of *A3* genes counted in this study might underestimate from the true value because of relatively low resolution of many whole genome sequences. Moreover, we particularly focused on the numbers and sequences of the Z domain of *AID/APOBEC* family genes, and we could not fully address whether (1) some two Z domains compose a double domain gene and (2) there are splicing variants. Nevertheless, here we covered most mammalian linages and showed a clear correlation between the number of ERVs and the number of *A3* genes. Moreover, the greater the number of *A3* genes encoded by a given host, the more G-to-A mutations are detected in their ERVs. To our knowledge, this is the first investigation comprehensively describing the evolutionary relationship between ancient retroviruses and *A3* family genes in a broad range of mammalian species.

## Methods

### Identification of sequences related to *AID/APOBEC* family genes in mammalian genomes

Information of the genome sequences analyzed in this study is summarized in **Supplementary Table 1**. To retrieve sequences homologous to *AID/APOBEC* family genes, we first screened these genomes using The DIGS tool (http://giffordlabcvr.github.io/DIGS-tool/), a tBLASTn-based method. As the queries, amino acid sequences of *AID/APOBEC* family genes of five mammals (human, mouse, cow, megabat, and cat) were used (**Supplementary Fig. 1** and **Supplementary Table 2**). Threshold of the bit score in the tBLASTn search was set at 50 bit. Of the hit sequences in the DIGS search, we subsequently extracted the regions including the sequence corresponding to the conserved sequence of *AID/APOBEC* family genes^24^ (“extracted region” is summarized in **Supplementary Fig. 1a**). Since this conserved sequence is located on a single exon in *AID/APOBEC* family genes (**Supplementary Fig. 1c**), this sequence can be extracted continuously. Of these hit sequences, we removed a sequence in which the sequence alignment does not cover >70% of the conserved sequence in the query (**Supplementary Fig. 13**). We also discarded hit sequences containing undetermined nucleotides (i.e., Ns). To further remove unreliable hit sequences, we performed a phylogenetic tree-based filtering. First, we constructed the phylogenetic tree of the hit sequences using neighbor-joining (NJ) method^66^ implemented in MEGAX^67^. Subsequently, the outlier sequence, which has >5 Z score with respect to the external branch length in the tree, was detected and discarded. The final set of the identified sequences of *AID/APOBEC* family genes is summarized in **Supplementary Table 3**.

Phylogenetic classification of the identified sequences of *AID/APOBEC* family genes was performed according to the following procedures: multiple sequence alignment (MSA) of the sequences was constructed using MAFFT (version 7.407)^68^ with the algorism L-INS-I. Phylogenetic tree was reconstructed using NJ method^66^ implemented in MEGAX^67^. Alignment sites with the >85% site coverages were used for the tree construction. According to the tree topology, we categorized the identified sequences into nine groups as summarized in **Supplementary Table 3**.

The identified sequences of *AID/APOBEC* family genes were classified into intact or pseudogenized genes according to the presence or absence of the premature stop codons. If a sequence contains ambiguous nucleotides (e.g., R or Y), the sequence was assigned the category “not determined (in **Fig. 1b**)”.

### Characterization of evolutionary features of *AID/APOBEC* family genes

For each class of *AID/APOBEC* family genes, MSA of the nucleotide sequences was constructed using MUSCLE^69^ with the codon-based alignment algorism. Codon sites with >50% site coverages were only used for the downstream analyses. The MSA was translated to the amino acid MSA. Logo plots of the amino acid sequences were generated using weblogo3^70^. Positional Shannon’s entropy score of the amino acid sequences was calculated using an web application in the Los Alamos HIV-1 sequence database (www.hiv.lanl.gov/content/sequence/ENTROPY/entropy_one.html). To detect codon sites under the positive or diversifying selection in *AID/APOBEC* family genes, dN/dS ratio test with branch-site model was performed using Hyphy MEME^28^. Phylogenetic tree used in this test was constructed by maximum likelihood method implemented in MEGAX^67^.

### Characterization of the genomic location of *AID/APOBEC* family genes

To determine the canonical genomic locus of *A3* genes (i.e., the region sandwiched by *CBX6* and *CBX7*), the genomic locations of *CBX6* and *CBX7* were determined using The DIGS tool (http://giffordlabcvr.github.io/DIGS-tool/). The threshold of tBLASTn search in DIGS analysis was set at 50 bit. We only analyzed a genome in which *CBX6* and *CBX7* were detected on the same scaffold in the downstream analysis. Sequences of *AID/APOBEC* family genes identified in this study were classified with respect to the presence or absence in the canonical *A3* locus (**Supplementary Table 4**). Genomic synteny of *AID/APOBEC* family genes was illustrated using ggplot2 (https://ggplot2.tidyverse.org/) with the library ggquiver (https://github.com/mitchelloharawild/ggquiver) implemented in R.

To characterize *A3* genes in the outside of the canonical *A3* locus, the gene structures of the outside *A3* genes in two New World monkeys (*Saimiri boliviensis* and *Aotus nancymaae*) were examined. To extract the full-length sequence of the non-canonical (i.e., “outside”) *A3* genes, genome sequences of these two species were scanned by BLASTn search using the query sequence of human *A3G* mRNA (Genbank accession number NM_021822.4). Information of the outside *A3* genes identified in this study is summarized in **Supplementary Table 5**. The gene structures were illustrated using the low-level plotting functions pre-implemented in R.

### RNA-Seq analysis of *AID/APOBEC* family genes

To detect expression of *AID/APOBEC* family genes in New World monkey tissues, RNA-Seq data in public databases was analyzed. RNA-Seq dataset used in the present study is summarized in **Supplementary Table 6**. RNA-Seq reads were trimmed by Trimmomatic (version 0.36)^71^ and subsequently mapped to the reference genomes using STAR (version 020201)^72^. Reads mapped on the identified loci of *AID/APOBEC* family genes were counted using featureCounts (version 1.6.4)^73^. In the read count, a read mapped to a unique genomic region was only counted. Read counts were normalized with the total uniquely mapped reads, and expression levels were measured as fragments per kilobase per million mapped fragments (FPKM).

### Extraction of the repetitive sequences derived from TEs in mammalian genomes

Sequences of TEs were extracted using RepeatMasker (version open-4-0-9) (http://repeatmasker.org) with Repbase RepeatMasker libraries (version 20181026)^74^. As the search engine, RMBlast was selected. RepeatMsker was run with options “-q xsmall -a -species <species>” where <species> is the species name of the analyzed genome. Information about the repetitive sequence compositions in the genomes is summarized in **Supplementary Table 7**.

### Detection of footprints of A3 activity in ERVs

To assess the accumulation level of G-to-A mutations in ERVs and the other TEs, the strand bias of the G-to-A mutation rate was calculated as the following procedures: First, mutations were detected in each TE integrant compared to the consensus sequence of the TE group. For this comparison, the pairwise sequence alignment outputted by RepeatMasker was used. TE integrant with low-confidential alignments (<1000 Smith-Waterman score) was excluded from the analysis. Second, G-to-A mutation rates in the positive and negative strands were calculated. Finally, the strand bias score was defined as a rate ratio of the G-to-A mutation between the positive and negative strands (i.e., the mutation rate in the positive strand was divided by the one in the negative strand). The strand bias score was calculated for each TE integrant or each TE group. Statistical significance of the strand bias was evaluated by Fisher’s exact test with false discovery rate calculation by Benjamini-Hochberg method.

Site preference of the G-to-A mutations in human ERVs was evaluated at the dinucleotide context. This preference was assessed for each group of ERVs. G-to-A mutation rates in <u>G</u>A, <u>G</u>T, <u>G</u>C, and <u>G</u>G sites were separately calculated, and the variation of the mutation rates among these four sites was evaluated as coefficient of variation (i.e., standard deviation divided by mean).

### Estimation of insertion dates of ERVs

Insertion dates of respective ERV loci were estimated by the two distinct methods: the ortholog distribution-based and genetic distance-based methods. The ortholog distribution-based estimation was performed for ERVs in human and mouse genomes. Liftover chain files were downloaded from UCSC genome browser (https://genome.ucsc.edu/) (**Supplementary Table 8**). Using Liftover (http://genome.ucsc.edu/cgi-bin/hgLiftOver) and the chain file, the genomic coordinate of ERV integrant in one genome was tried to be converted to that in another genome with the option “minMatch=0.5”. If this conversion succeeded, we judged that the orthologous copy of the ERV integrant was present in the corresponding genome. In case of mouse ERVs, we first converted genomic coordinates of ERVs in “Mm9” to those in “Mm10”, which is the latest version of the mouse reference genome. Subsequently, the genomic coordinates in “Mm10” converted to those in the genomes of respective species. Insertion dates of ERVs were estimated from the ortholog distributions according to the scheme summarized in **Supplementary Fig. 14**.

The genetic distance-based estimation of insertion dates was performed for ERVs in Primates, Rodents, Insectivora, and Marsupialia. Genetic distance of each ERV integrant from the consensus sequence was calculated using RepeatMasker (http://repeatmasker.org). Subsequently, the amounts of ERV insertions in respective genetic distances were summarized using the function “Repeat Landscape” implemented in RepeatMasker. The genetic distance was converted to the insertion date under the molecular clock assumption. As the mutation rate for Primates, Insectivora, and Marsupialia, 2.2 × 10^−9^ mutation/year/site^75^ was used. Under this mutation rate, the estimated insertion dates were highly concordant between the genetic distance-based and ortholog distribution-based methods for human ERVs (**Supplementary Fig. 15**). As the mutation rate for Rodents, 7.0 × 10^−9^ mutation/year/site was used. This is because (1) evolutionary rate in Rodents is known to be much faster than those in the other mammals, particularly after the divergence between mouse and rat^76^; and (2) under this mutation rate, the estimated insertion dates were highly concordant between the genetic distance-based and ortholog distribution-based methods for mouse ERVs (**Supplementary Fig. 16**).

## Acknowledgments

We would like to thank Mai Suganami (Division of Systems Virology, Institute of Medical Science, The University of Tokyo, Japan) for technical support.

This study was supported in part by AMED J-PRIDE 19fm0208006s0103 19fm0208006h0003 (K.S.); AMED Research Program on HIV/AIDS 19fk0410014h0002 (to K.S.) and 19fk0410019h0002 (to K.S.); JST CREST (to K.S.); Grants-in-Aid for Scientific Research (KAKENHI) Scientific Research B 15H05707 18H02662 (to K.S.), Scientific Research on Innovative Areas 16H06429 (to K.S.), 16K21723 (to K.S.), 17H05813 (to K.S.), and 19H04826 (to K.S.), and Fund for the Promotion of Joint International Research (Fostering Joint International Research) 18KK0447 (to K.S.); JSPS Research Fellow PD 19J01713 (to J.I.); Takeda Science Foundation (to K.S.); ONO Medical Research Foundation (to K.S.); Ichiro Kanehara Foundation (to K.S.); Lotte Foundation (to K.S.); Joint Usage/Research Center program of Institute for Frontier Life and Medical Sciences, Kyoto University (to K.S.); and JSPS Core-to-Core program (A. Advanced Research Networks) (to R.J.G. and K.S.). R.J.G. was supported by a grant from the UK Medical Research Council (No. MC_UU_12014/10).

## Author contributions

J.I. mainly performed investigations; J.I. and K.S. prepared the figures; all authors contributed to data interpretation, designed the research, wrote the paper, and approved the final manuscript.

