## Supplementary Figures 1-16 for "Rapid evolution of antiviral *APOBEC3* genes driven by the conflicts with ancient retroviruses"

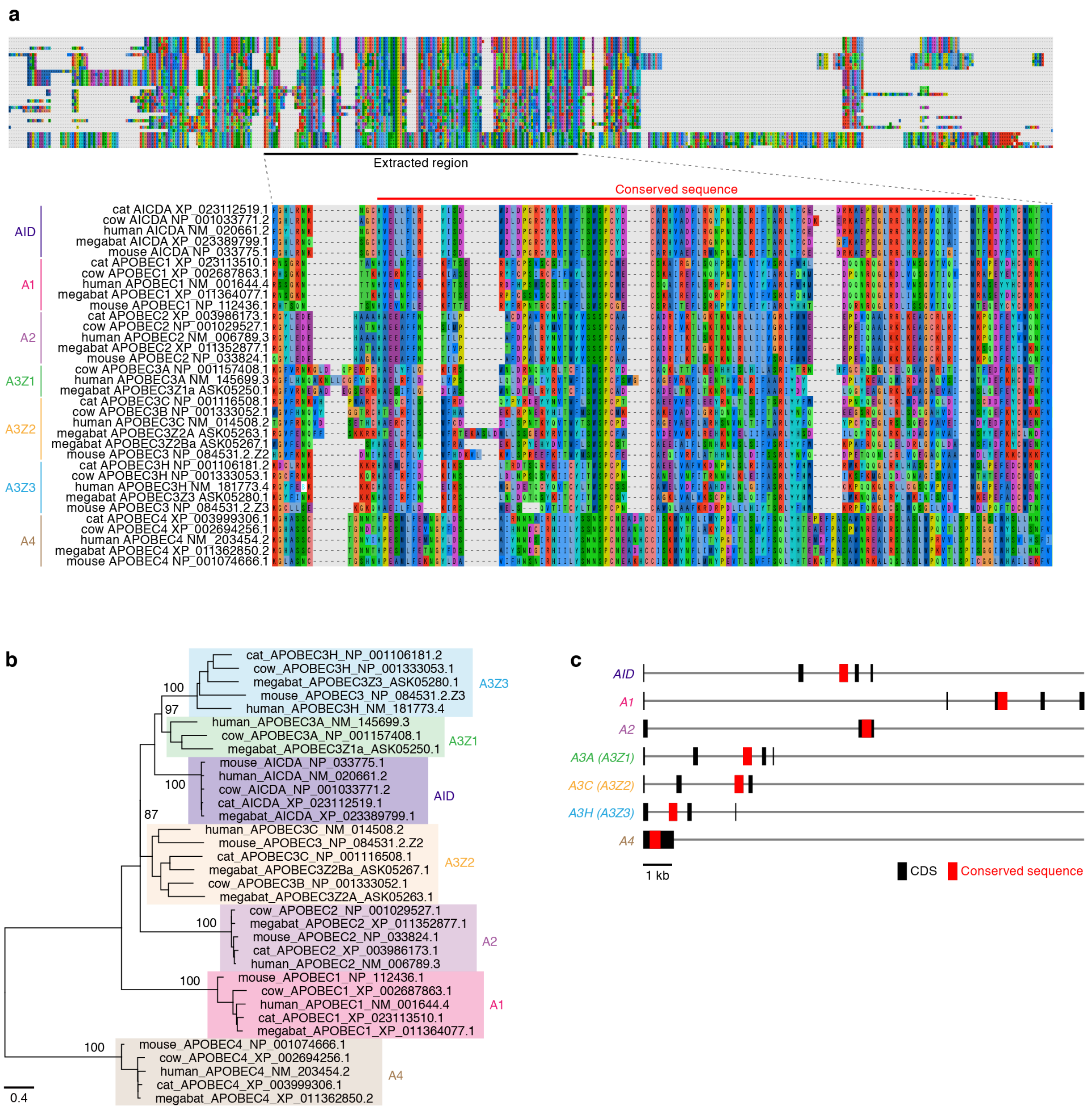


**Supplementary Fig. 1. Conserved sequence of *AID/APOBEC* family genes.**
**a** Amino acid sequences of *AID/APOBEC* family genes that were used as queries in the DIGS (tBLASTn) search. Of the hit sequences in the DIGS search, the regions including the sequence corresponding to the conserved sequence of *AID/APOBEC* family genes were extracted (referred to as “Extracted region”).
**b** Phylogenetic tree of the conserved sequences of *AID/APOBEC* family genes. The tree was constructed by maximum likelihood (ML) method. Bootstrap values are indicated at the nodes.
**c** Gene structure of *AID/APOBEC* family genes in the human genome. Black and red boxes indicate the regions corresponding to the coding sequence (CDS) and the conserved sequence, respectively. Note that the conserved sequence is located on a single exon in all types of *AID/APOBEC* family genes.

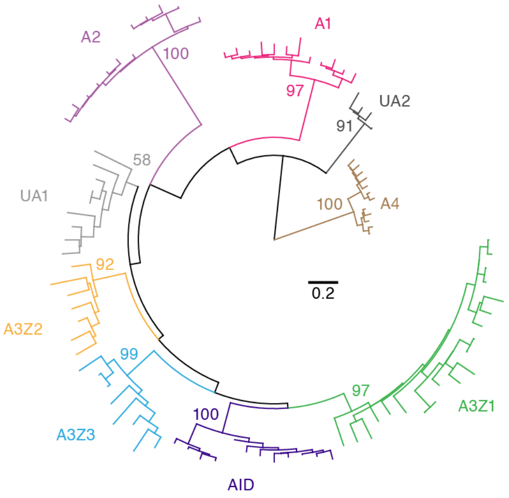


**Supplementary Fig. 2. Phylogenetic tree of *AID/APOBEC* Z domains identified in Afrotheria, Xenarthra, and Marsupialia genomes.**
The tree was reconstructed by maximum likelihood (ML) method based on the nucleic acid sequences. Bootstrap values are indicated at the nodes.

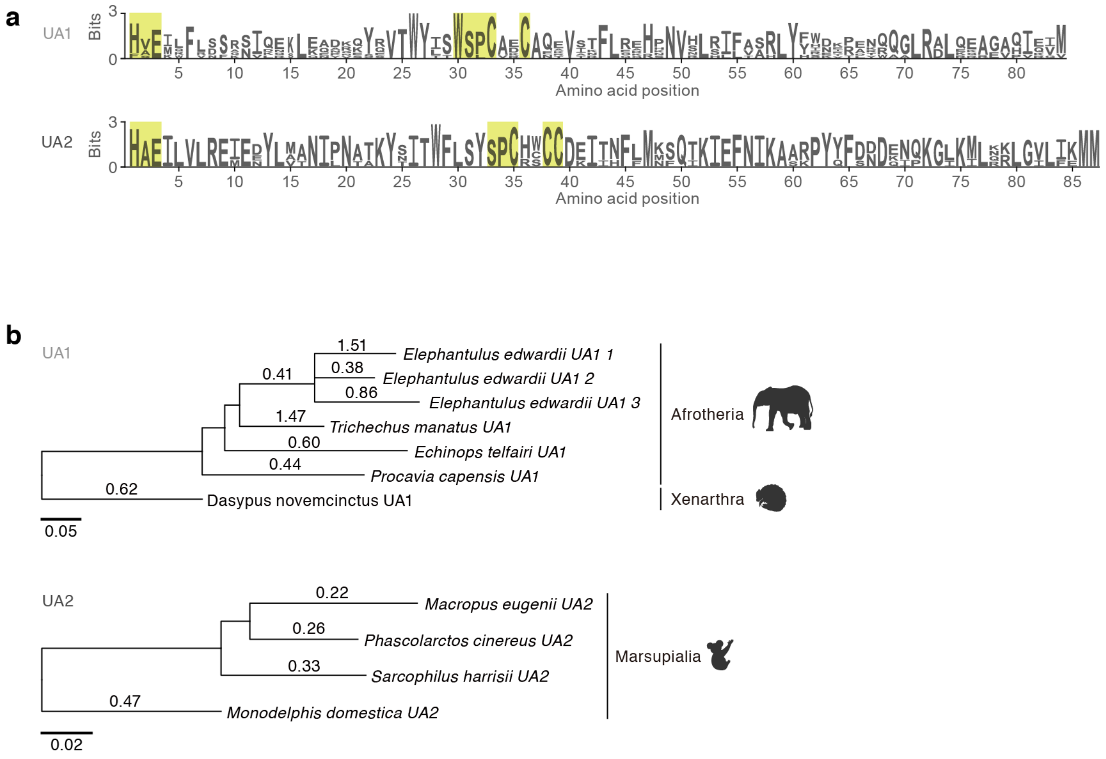


**Supplementary Fig. 3. Characterization of the lineage-specific *AID/APOBEC* genes (*UA1* and *UA2* genes).**

**a** Logo plots of amino acid sequences of UA1 and UA2. Yellow squares indicate the amino acid residues corresponding to the catalytic domain of AID/APOBEC proteins.

**b** Phylogenetic tree of *UA1* and *UA2* Z domains. The tree was reconstructed by maximum likelihood (ML) method based on the nucleic acid sequences. Estimated dN/dS ratios are indicated on the branches. Note that purifying selections (dN/dS < 1) on *UA1* and *UA2* Z domains were detected.

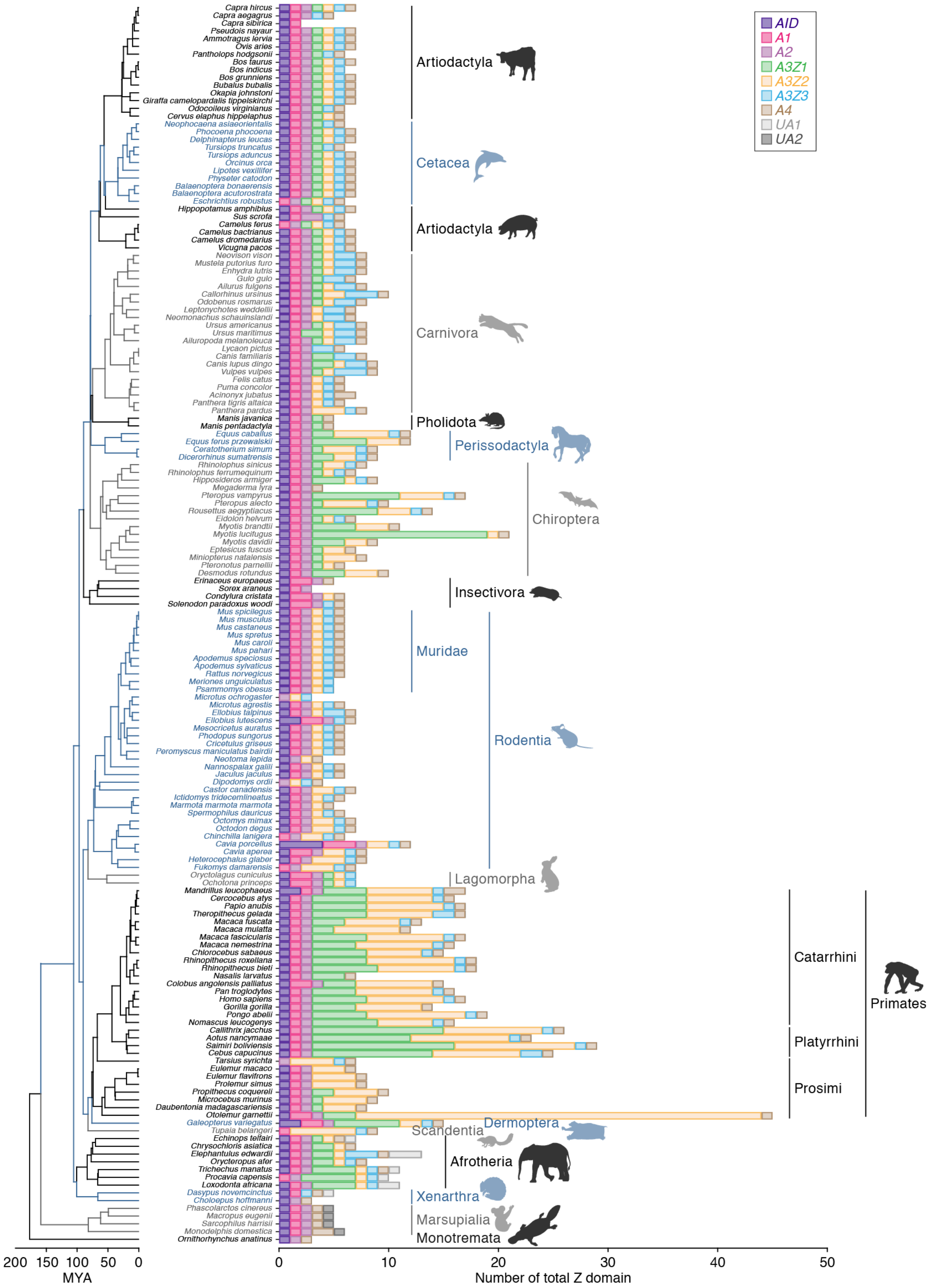


**Supplementary Fig. 4. Number of the identified *AID/APOBEC* Z domains in each mammal species.**

Unlike **Fig. 2**, both of intact and pseudogenized *AID/APOBEC* Z domains were counted.

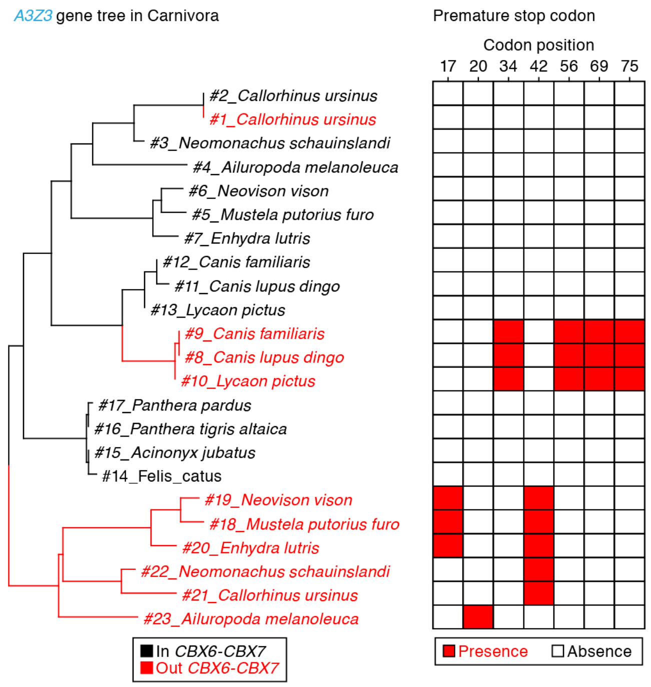


**Supplementary Fig. 5. Gene duplications of *A3Z3* followed by pseudogenizations in Carnivora.**

(Left) Phylogenetic tree of *A3Z3* in Carnivora. Red and black respectively indicate *A3Z3* located in the inside and outside of the canonical *A3* gene locus, sandwiched by *CBX6* and *CBX7* genes.

(Right) Presence of premature stop codons. Positions of the stop codons in the multiple sequence alignment are indicated.

Note that gene duplications of *A3Z3* followed by pseudogenizations occurred at least twice in Carnivora.

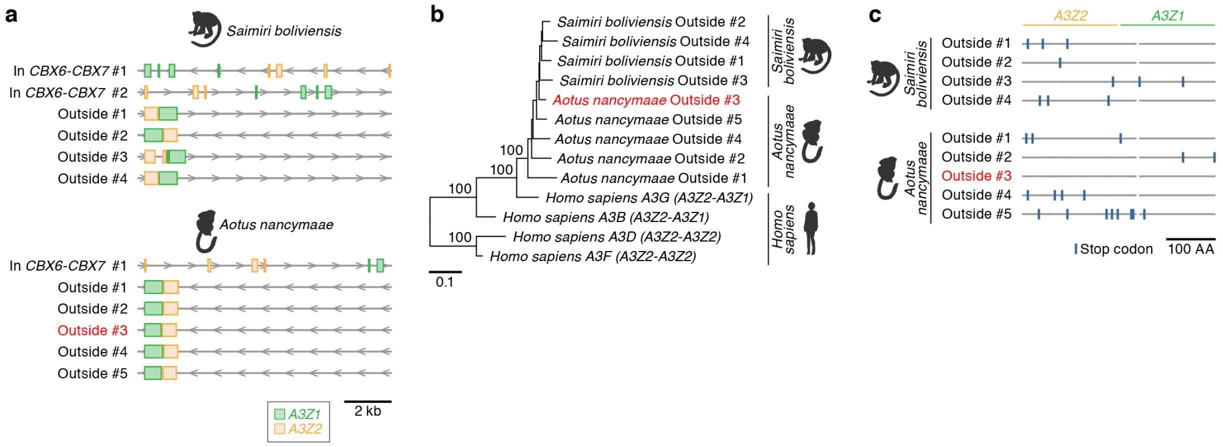


**Supplementary Fig. 6. Retrotransposition of *A3G*-like genes in New World monkeys.**
**a** Gene structures of *A3Z2-A3Z1* type genes in the genomes of New World monkeys. Genes within the canonical *A3* gene locus ("in *CBX6-CBX7*") and the ones outside the locus ("outside") are shown. The arrowheads indicate the direction of respective loci.
**b** Phylogenetic tree of the retrotransposed *A3Z2-A3Z1* type genes in the two New World monkeys and human double domain *A3* genes. Note that the retrotransposed genes formed a cluster with human *A3G* (*A3Z2-A3Z1* type) gene.
**c** Presence or absence of premature stop codons in the retrotransposed *A3Z2-A3Z1* type genes in the two New World monkeys.

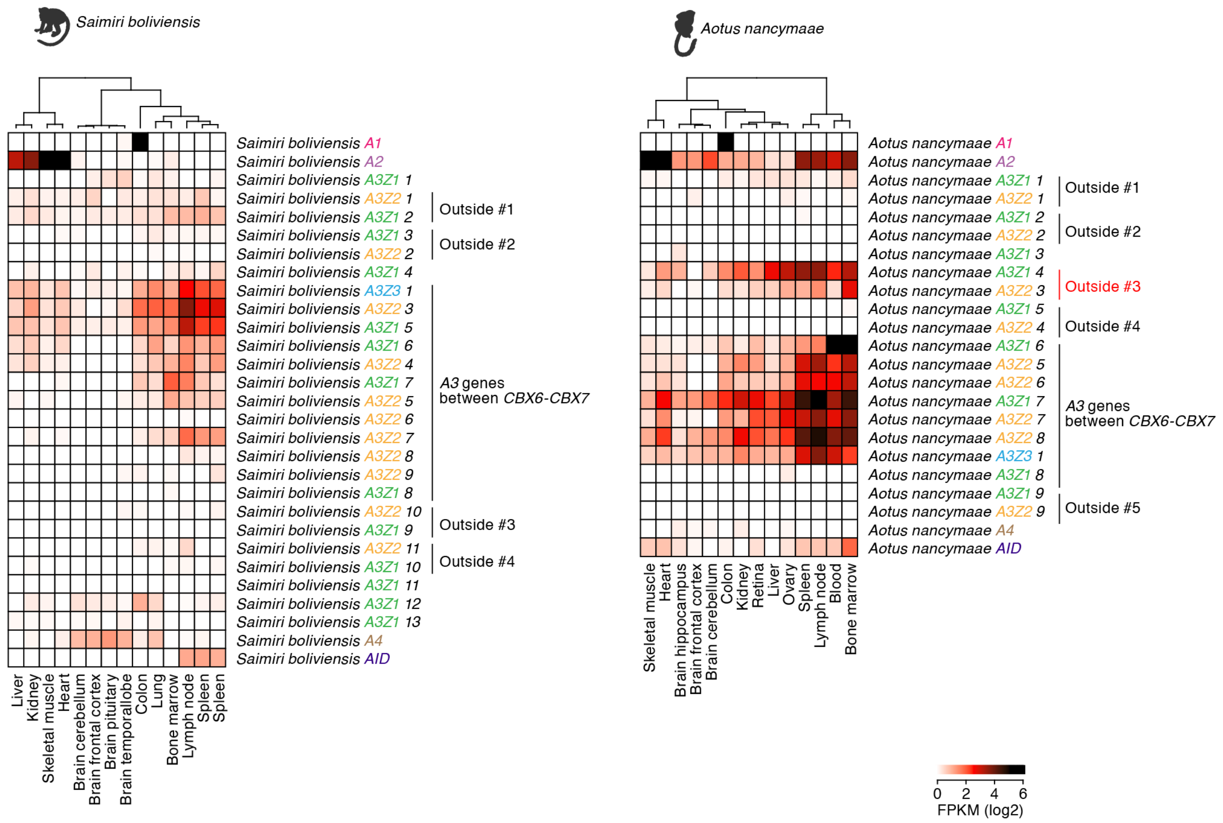


**Supplementary Fig. 7. RNA expression of the retrotransposed *A3* genes in the tissues of New World monkeys.**
RNA-Seq data in public databases were analyzed. Rows indicate the identified sequences of *AID/APOBEC* family genes, and columns indicate the tissues of New World monkeys. Color indicates RNA expression level (log2 FPKM). Note that the "outside #1-4" genes in *Saimiri boliviensis* and "outside #1-5" genes in *Aotus nancymaae* are identical to those in **Supplementary Fig. 6.**

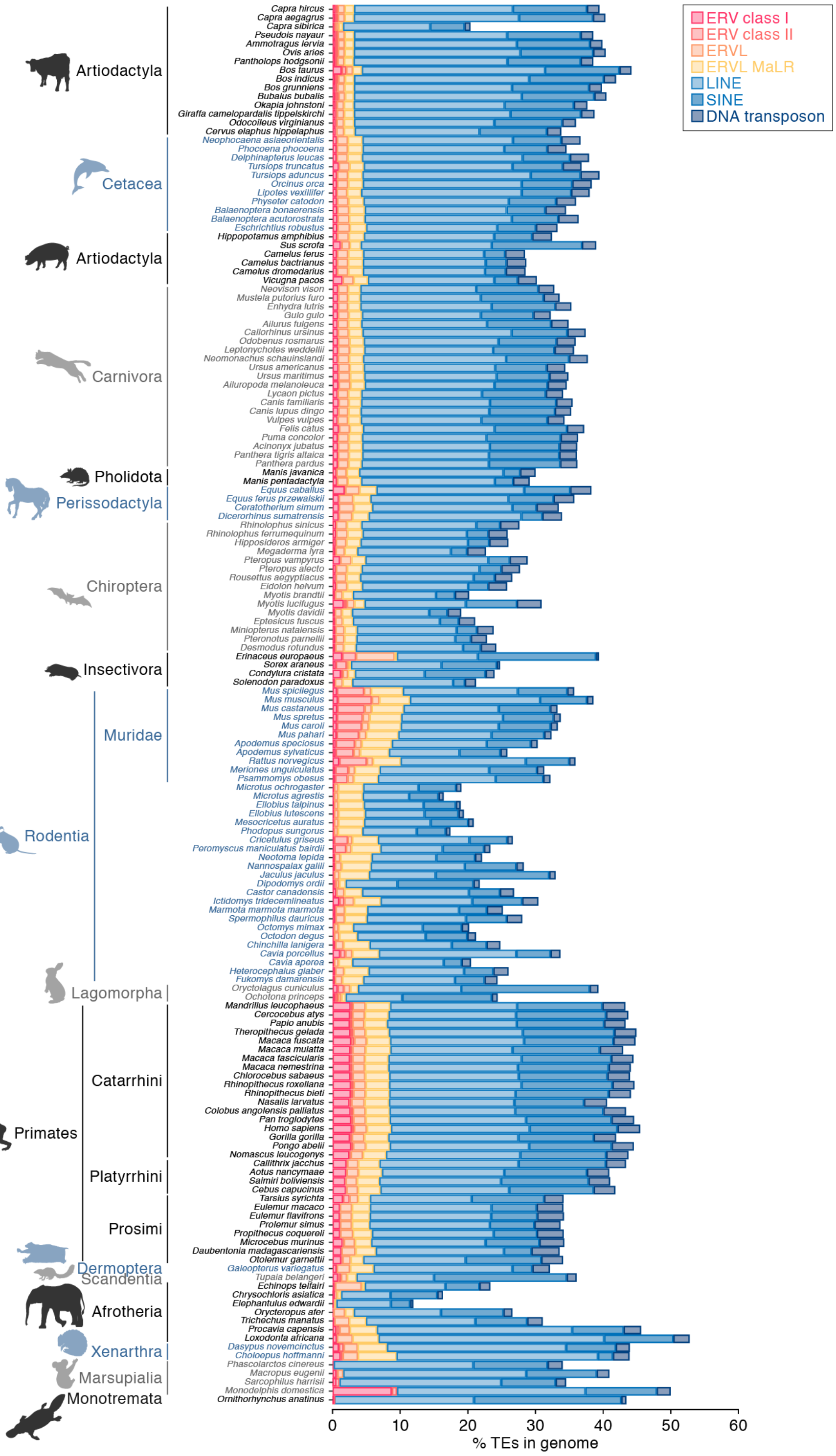


**Supplementary Fig. 8. Proportions of TE sequences in the genomes of respective mammalian species.**

Proportions of ERV sequences in the genomes of respective mammalian species are also shown in **Fig. 4a**.

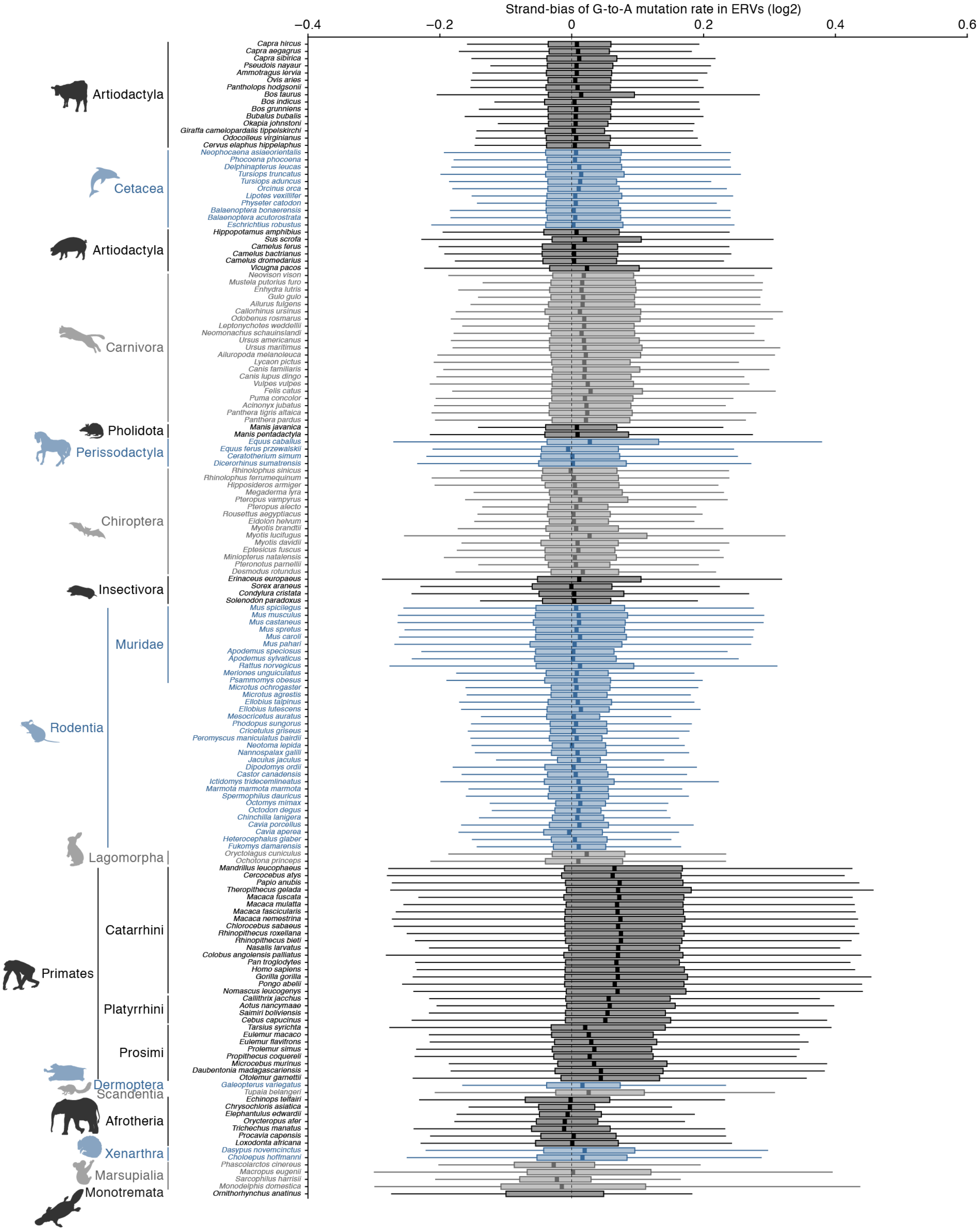


**Supplementary Fig. 9. Difference in the strand biases of G-to-A mutation rates in ERVs among mammals.**

Log2-transformed strand bias score of G-to-A mutations (i.e., the mutation rate in the positive strand divided by the one in the negative strand) is shown.

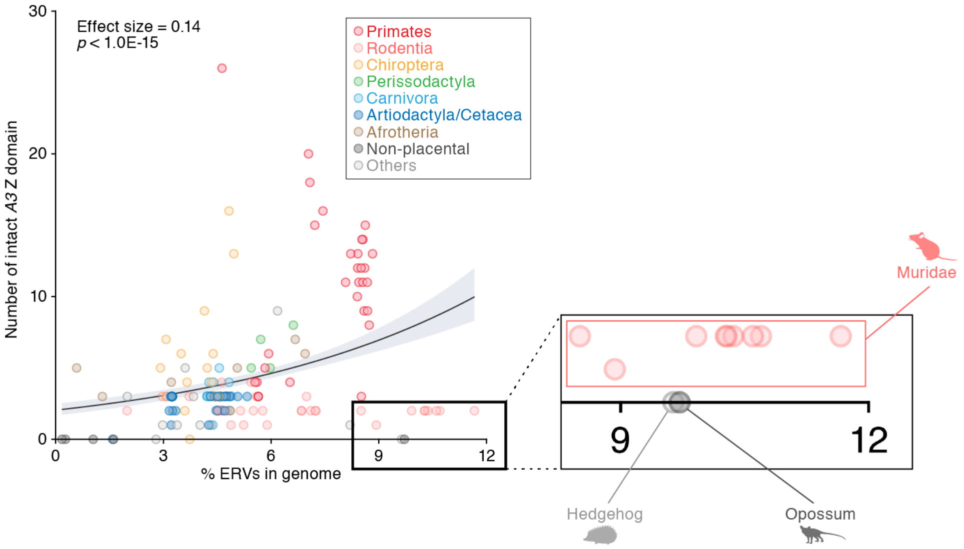


**Supplementary Fig. 10. Outlier species, which were with higher ERV invasions but possessed few or no *A3* genes.**
The figure is identical to **Fig. 5a**. To indicate the outlier species (Muridae, hedgehog, and opossum), the dots of these species are zoomed up.

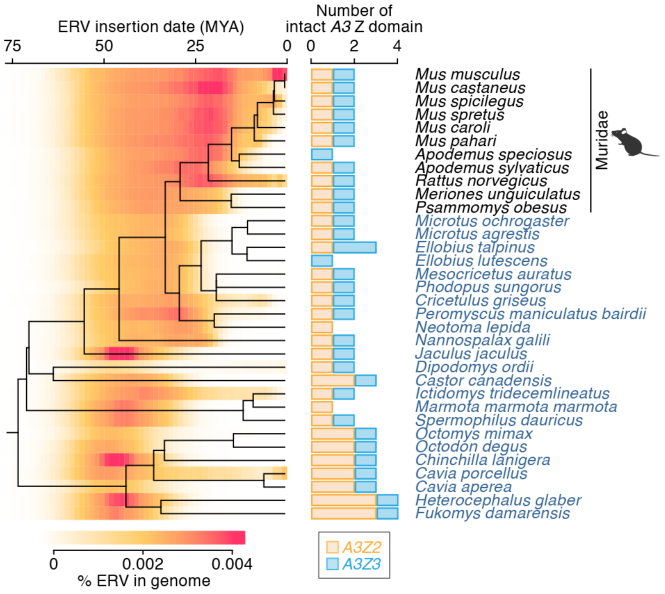


**Supplementary Fig. 11. Insertion dates of ERVs and the numbers of *A3* genes in Rodents.**

(Left) Amount of ERV insertions at each age in respective rodent species. ERV insertion date was estimated according to the genetic distance of each ERV integrant from the consensus sequence under the molecular clock assumption (7 × $10^{-9}$ mutation/site/year). (Right) Number of intact *A3* Z domains.

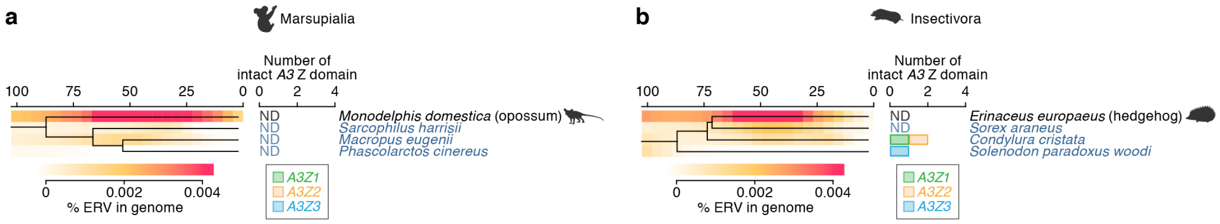


**Supplementary Fig. 12. Insertion dates of ERVs and the numbers of *A3* genes in Marsupialia and Insectivora.**

(Left) Amount of ERV insertions at each age in respective animal species. ERV insertion date was estimated according to the genetic distance of each ERV integrant from the consensus sequence under the molecular clock assumption ($2.2 \times 10^{-9}$ mutation/site/year). (Right) Number of intact *A3* Z domains.

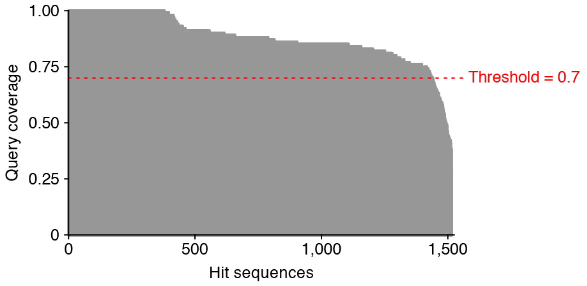


**Supplementary Fig. 13. Coverages of sequence alignment in query sequences in the DIGS (tBLASTn) search.**

Y-axis indicates the proportion of the length of the query sequence covered by the alignment (referred to as coverage). Of these hit sequences in the DIGS search, a sequence with <70% of the coverage was discarded.

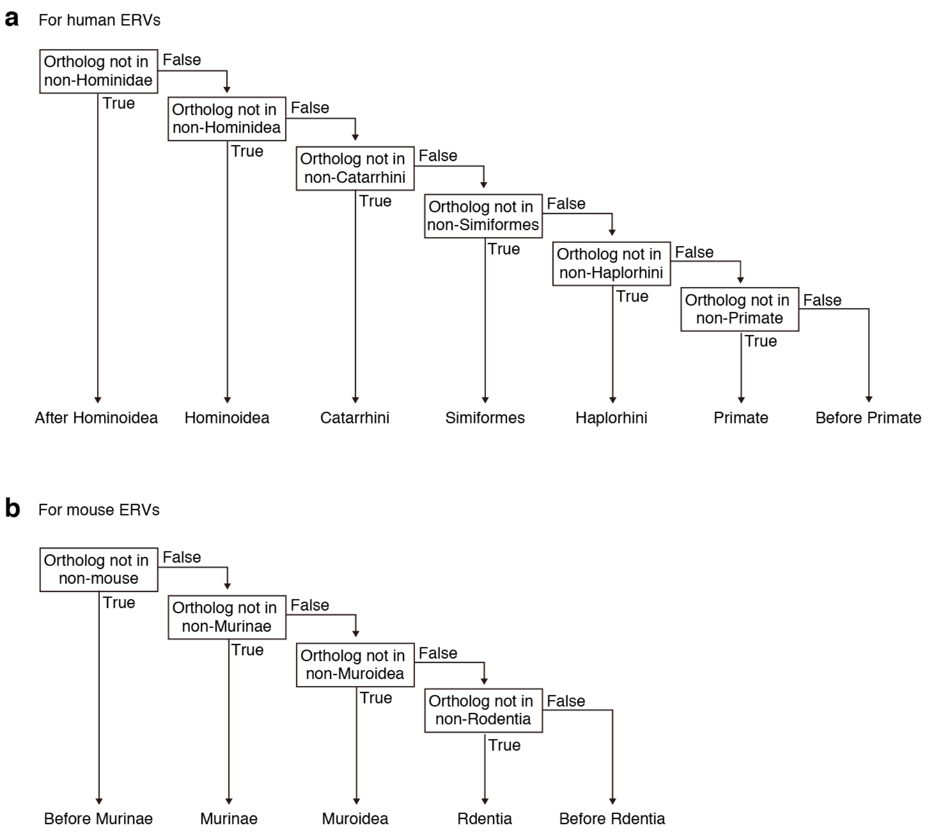


**Supplementary Fig. 14. Flow chart to assign the insertion dates of ERVs according to the ortholog distribution.**

The flow chart for ERVs in the genomes of human (**a**) and mouse (**b**) are shown.

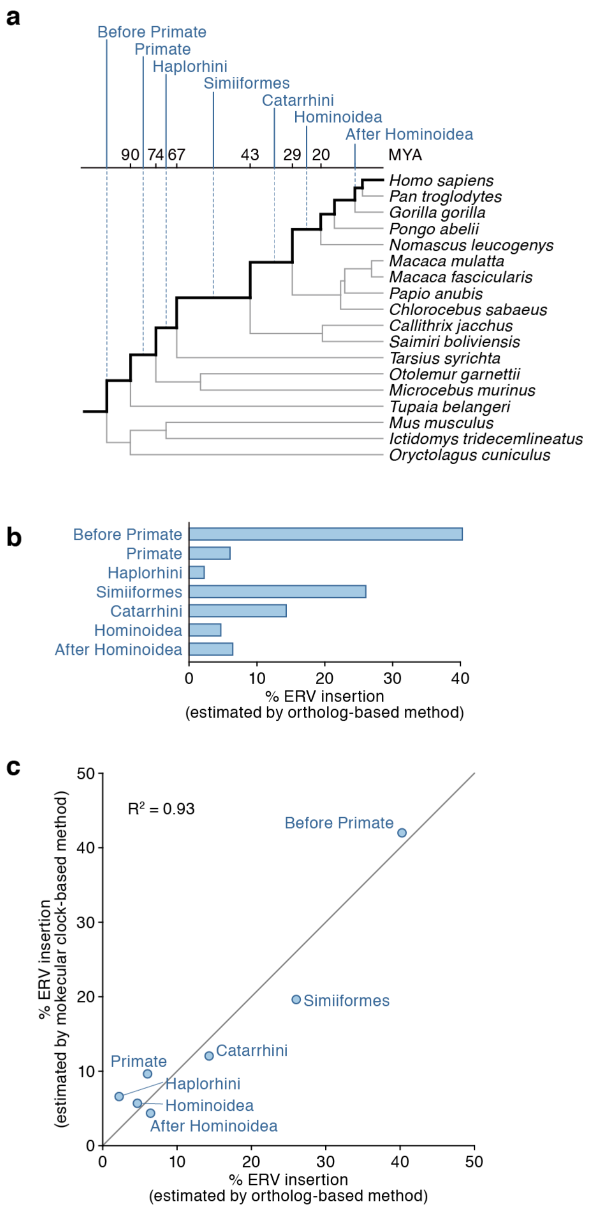


**Supplementary Fig. 15. Comparison of ERV insertion dates estimated by the genetic distance-based and ortholog distribution-based methods in human ERVs.**

**a** Stratification of human ERVs according to the estimated insertion dates.

**b** Amounts of ERV insertions in respective periods estimated by the ortholog distribution-based methods.

**c** Comparison of ERV insertion amounts estimated by the two methods. In the genetic distance-based method, $2.2 \times 10^{-9}$ mutation/site/year was used as the mutation rate. The line y = x and coefficient of determination ($R^2$) are shown.

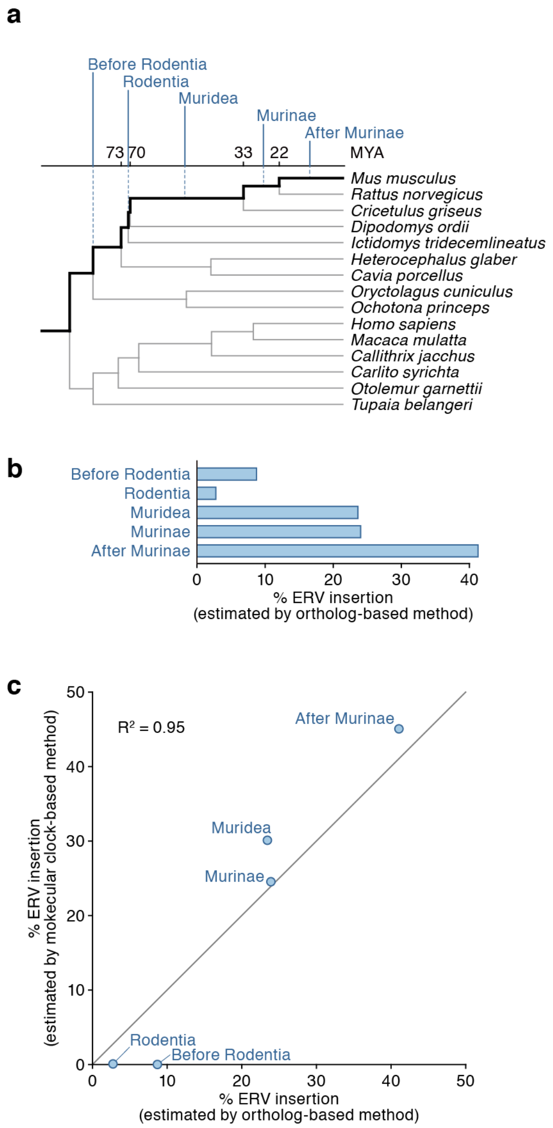


**Supplementary Fig. 16. Comparison of ERV insertion dates estimated by the genetic distance-based and ortholog distribution-based methods in mouse ERVs.**

**a** Stratification of mouse ERVs according to the estimated insertion dates.

**b** Amounts of ERV insertions in respective periods estimated by the ortholog distribution-based methods.

**c** Comparison of ERV insertion amounts estimated by the two methods. In the genetic distance-based method, $7 \times 10^{-9}$ mutation/site/year was used as the mutation rate. The line y = x and coefficient of determination ($R^2$) are shown.
